# Sero-prevalence of brucellosis, Q-fever and Rift Valley Fever in humans and livestock in Somali region, Ethiopia

**DOI:** 10.1101/2020.01.31.928374

**Authors:** Mohammed Ibrahim, Esther Schelling, Jakob Zinsstag, Jan Hattendorf, Emawayish Andargie, Rea Tschopp

## Abstract

Information on zoonotic diseases in humans and livestock are limited in pastoral/agro-pastoral communities in Ethiopia. A multi-stage cross sectional cluster design study was implemented with the aim to establish the seroprevalence of zoonotic diseases including brucellosis, Q-fever and Rift Valley Fever (RVF) in humans and livestock in Adadle woreda of the Somali region, Ethiopia. Blood samples were collected from humans and livestock and tested by relevant serological tests. For brucellosis, Rose Bengal test (RBT) and indirect ELISA was used for screening and confirmatory diagnosis respectively. Indirect and competitive ELISA were also used for Q-fever and RVF respectively. The individual seropositivity of Q-fever in livestock was 9.6% (95% CI 5.9-15.1) in cattle, 55.7% (95% CI 46.0-65.0) in camels, 48.8% (95% CI 42.5-55.0) in goats, and 28.9% (95% CI 25.0-33.2) in sheep. In humans, seropositivity of Q-fever was 27.0% (95% CI 20.4-34.0), with prevalence in males of 28.9% vs 24.2% in females (OR= 1.3; 95% CI 0.6-2.5). Camel seropositivity of Q-fever was significantly associated with age (OR= 8.1; 95% CI 2.8-23.7). The individual apparent seroprevalence of RVF was 13.2% (95% CI 8.7-18.8) in humans, 17.9 % (95% CI 11.0-27.8) in cattle, 42.6% (95% CI 34.8-50.7) in camels, 6.3% (95% CI 3.3-11.6) in goats and 7.4% (95% CI 4.7-11.5) in sheep. Camels had the highest seropositivity of both Q-fever (55.7%; 95% CI 46.0-65.0) and RVF (42.6%; 95% CI 34.8-50.7). Only a weak correlation was observed between human and livestock seropositivity for both Q-fever and RVF. Only cattle and camels were seropositive for brucellosis by iELISA. The individual seroprevalence of brucellosis was 2.8(0.9-6.4) in humans, 1.5% (95% CI 0.2-5.2) in cattle and 0.6% (95% CI 0.0-3.2) in camels. This study showed the importance of zoonoses in Somali regional state and is the first published study to describe RVF exposure in humans and livestock in the country. Collaboration between public and animal health sectors for further investigation on these zoonoses using the One Health concept is indispensable.

## 1. Introduction

Zoonoses are infectious diseases transmitted between human and vertebrate animals. These diseases include those from animal sources food. The international communities do not address neglected zoonotic diseases (NZDs) adequately [1]. Brucellosis, Q-fever and Rift Valley Fever are among those NZDs, which are largely eliminated in developed countries but under-diagnosed and under-reported in developing countries [2]. Effective management of zoonoses benefits from a One Health approach, creating synergistic benefits from the collaboration of human and animal health sectors [3]. Ethiopia is among the top five countries with the highest zoonotic infections in the world [4]. Despite its burden, attention by the government rose only recently, where the five most prevalent zoonotic diseases were prioritized as following: Rabies, anthrax, brucellosis, leptospirosis and echinococcosis [5].

Brucellosis is one of the neglected bacterial zoonoses, which have economic importance globally [6]. This disease is caused by the genus *Brucella*. The economically most important species are *B. melitensis* and *B. abortus* having a high potential of human infection [3] affecting small ruminants and cattle respectively [7]. Transmission from animals to humans occurs usually due to consumption of unpasteurized milk and milk products or direct contact with infected animal especially during parturition, with direct contact with placentas or aborted fetuses [8]. Human brucellosis causes a flu-like illness with a fever, weakness, malaise, myalgia and weight loss. It can be debilitating in chronic stages with serious complications (e.g. endocarditis, musculoskeletal lesions) which can be potentially fatal if not treated. In livestock, *Brucella* spp cause abortion, infertility, and consequently, reduction of milk yields [7]. Human brucellosis infection shows non-specific symptoms and remains generally unnoticed or undiagnosed by medical doctors due to overlapping with other febrile illnesses [9]. Brucellosis occurs globally with high incidences in the Middle East [10].

In Ethiopia, livestock brucellosis is endemic and was reported in different studies [11–15]. Most studies were done in the highlands targeting urban and peri-urban dairy farms. Seroprevalence of cattle in extensive production systems is lower than that of intensive production systems [16]. The highest prevalence of brucellosis was recorded in central Ethiopia followed by the southern part, whereby lower prevalences were seen in the western and eastern parts. Camel seropositivity for brucellosis in Ethiopia ranged from 0.7 to 12% for the Rose Bengal Plate Test (RBPT) and 0.5 −10% for Complement Fixation Test (CFT) in different agro-ecologies [14]. Studies on human brucellosis in Ethiopia are sparse with less information about risk factors for human infection [13, 17].

Q-fever is a zoonotic disease caused by *Coxiella burnetii*, which is endemic worldwide except in New Zealand and Antaractica. It affects a wide range of mammals, birds and arthropods [18]. Domestic ruminants such as cattle, goats and sheep are the main reservoirs for Q-fever in humans [19]. Human infection occurs due to inhalation of dust contaminated by infected animal fluids, consumption of unpasteurized dairy products and contact with milk, urine, faeces, vaginal mucus or semen of infected animals. The most common sign of Q-fever in man is a flu-like illness, which can progress to an atypical pneumonia, resulting in a life threatening acute respiratory distress syndrome [20]. Infection in animals is predominantly asymptomatic but has been associated with late abortions, stillbirth, delivery of weak offspring and infertility [21].

Even though Q-fever have been given attention in developed countries, there are significant gaps in understanding the epidemiology of Q-fever infections in Africa [21]. Q-fever seropositivity among integrated human and animal studies was 13%, 23%, 33% and 16% in Egypt and 4%, 13%, 11% and 1% in Chad in cattle, goats, sheep and humans respectively [22, 23]. The seropositivity of Q-fever in camels was 80% in Chad and being a camel breeder was a risk factor of human seropositivity [23]. In Togo, people of Fulani ethnicity had greater livestock contact and a significantly higher seroprevalence than other ethnic groups (46% in Fulani vs 27% in non-Fulani) [20]. Reports of Q-fever sero-prevalence in various livestock species in Kenya, Ethiopia and Cote d’Ivoire varied between 9% and 90% while in humans it varied between 3% and 7% [2, 11, 21, 24].

Rift Valley Fever (RVF) is a peracute or acute zoonotic disease affecting ruminants and humans. It is caused by a mosquitoes borne virus of the Bunyaviridae family; genus *Phlebovirus* [25]. Rift Valley Fever epidemics in East Africa occur often when there is a heavy rainfall followed by flooding in arid and semi-arid areas favoring the massive hatching of mosquitoes eggs, whereof a part is already transovarially infected, and thus lead to rapid spread of the virus to animals and to a lesser extent to humans [26]. The majority of animal infections are due to bites of infected mosquitoes. In humans, RVF-Virus is transmitted by direct contact with infectious animal tissue or by the bites of infected mosquitoes [27]. The disease in ruminants and camels is characterized by abortion, neonatal mortality, weak-born offspring and liver damage in animals. In humans, most infections are asymptomatic or as a mild (flue-like) illness. In severe disease (about 7-8% of cases), it causes hemorrhage, encephalitis, visual disturbances and death [28].

Reports of RVF sero-prevalence in various livestock species in Kenya, Cote d’Ivoire, Chad, Tanzania and Western Sahara varied between 0% and 38% while in humans it was 0.8% [2, 29–32].

The ability of RVF to spread outside traditionally endemic countries, even out of the African continent lies in the fact that large ranges of arthropod vectors are capable of transmitting the virus. The presence of a wide range of hosts and vector species, and the epidemiological characteristics of RVF, had led to concerns that epidemics may occur in previously not described regions like Ethiopia [33]. In other East and central African countries such as Kenya, inter-epizootic/epidemic cases are increasingly documented for the past 10 years [34–37]. Ethiopia due to its geographic location as well as the vibrant livestock exchanges with neighboring countries makes it highly vulnerable to the disease particularly to cases that are not epidemic but occur on a more continued basis [38].

Somali region has the highest pastoralist communities in Ethiopia and yet, the status of the selected zoonotic diseases in humans and livestock are unknown. Thus, the aim of this study was to estimate the seroprevalence of brucellosis, Q-fever and RVF in humans and livestock and identify the associated risk factors in Adadle woreda. This study also aimed to highlight the awareness gap of the communities against zoonoses that could help shape future intervention strategies in preventing and controlling zoonotic diseases in the area.

## 2. Materials and methods

This study was part of research and development project called Jigjiga One Health Initiative (JOHI) funded by Swiss Agency for Cooperation and Development with major partnership between Jigjiga University, Swiss Tropical and Public Health Institute and Armauer Hansen Research Institute. The goal of the project was to improve the health and well-being of pastoralist communities in the Somali region of Ethiopia.

### 2.1. Study area

Adadle woreda (district) is situated in the Shabelle Zone of the Somali region of Ethiopia. It is located in the lowlands of the semi-arid Wabe Shabale River sub basin (Fig 1). The mean annual rainfall based on Gode (the main town of the zone) data is about 300 mm [39]. The main rainy season called “Gu” lasts from March to May and the short dry season known as “Xagaa” from June to August. The short rain “Dayr” between September and November and the long dry season “Jilaal” follow “Xagaa” from December to March. The woreda is composed of 15 kebeles (the smallest administrative units) [39] with a total population of 100,000 [40] (Fig 1). In 2000, the majority of people living in Adadle were pastoralists (60%), whereas 28% were agro-pastoralists and 10% practiced riverine cultivation as cited in [39].

**Fig 1.**
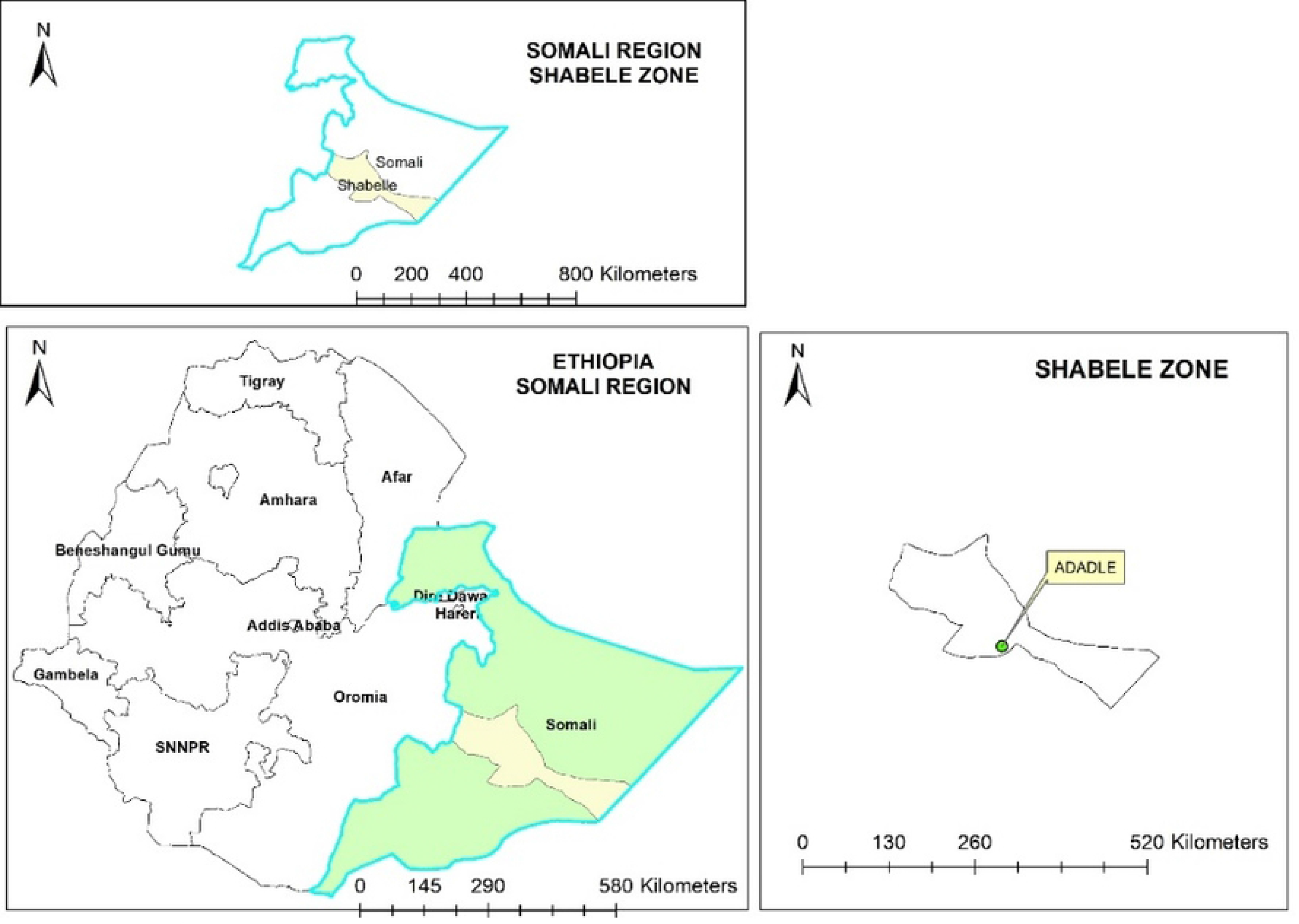
Map of the study area.

### 2.2. Sample size calculation

Sample size determination was conducted to estimate the precision of the study with an anticipated prevalence. In pastoral and settled livestock management systems in semi-arid areas of Africa, the seroprevalence of brucellosis in cattle is usually greater than 5%, ranging from 4.8-41.0% [41]. The seroprevalence of brucellosis is usually much lower in small ruminants than in cattle [41]. Considering that in the study area livestock has never been vaccinated against brucellosis, we assumed based on data from comparable countries that brucellosis had a prevalence of 7%, 5%, 12% and 7% in humans, camels, small ruminants and cattle, respectively. The design effect D was derived from the following formula D = 1 + (b-1) rho; where b is the number of units sampled per cluster and rho (ρ) is the intra-cluster correlation coefficient [42]. A rho value for zoonoses (and infectious diseases more generally) is usually between 0.05-0.2 and rarely exceeds 0.3 with highly contagious viral infections [42, 43]. Thus, a rho value of 0.15 was taken for initial sample size calculation. We calculated that a sample of 180 humans from 60 clusters will lead to a standard error of 2.2% of our estimate. Sampling of three hundred goats and three hundred sheep will lead to a standard error of 2.0% of our estimate for each species. Furthermore, sampling of 150 camels will lead to a standard error of 2.3% of our estimate.

### 2.3. Sampling procedure

Adadle woreda has 15 kebeles. Two kebeles were excluded from the study due to the lack of mobile phone network and poor accessibility. Six kebeles were selected randomly from the remaining thirteen kebeles with a selection probability proportional to the human population size. Melkasalah and Harsog were pure pastoralist kebeles, whereas Boholhagare, Bursaredo, Higlo and Gabal were agropastoralist. Even though Boholhagare and Higlo were listed as agropastoralist kebeles, people were mainly depended on livestock and practice crop plantation only during rainy seasons.

A village list was available for each agropastoral kebele. All villages in the kebele were assigned numbers. Community members (kebele administrators, elders and religious leaders) drew a minimum of 8 numbers from a bag to select the villages. In each selected village, households were selected by spinning a pen and proceeding in the direction of the pen head. All households in that direction were included. A village or camp was considered as a cluster in agropastoral or pastoral kebeles respectively. The two pastoralist kebeles were selected as follows: Kebele administrators reported which villages had concentrations of mobile pastoralist camps in the vicinity. We visited all reported villages and selected the camp (Reer) with the highest number of tents. We included all households of the selected Reer. Within the selected households, individuals who were present at the time of interview and were 16 years or older than were eligible to participate in the study.

#### 2.3.1. Livestock

The sampling was conducted between May and August, 2016 from six kebeles of Adadle woreda of Somali region, Ethiopia. The herd here is considered as a cluster. The animals within the herd of selected households were selected systematically using a sampling interval number (total number of animals in the herd which are ≥ 6 months divided by the number of animals to be sampled within the herd). The first animal was selected randomly, then every n^th^ animal until total sample size was attained. Camels were sampled outside the barn unlike other species but with the same methodology. Within each herd, a maximum of nine from each livestock species were sampled. A total of 171 camels, 297 goats, 269 sheep and 135 cattle were sampled from six kebeles.

#### 2.3.2. Humans

Individual people within the selected households whose animals were sampled who were ≥ 16 years and who provided informed consent to participate the study were sampled. Semi-structured questionnaires were conducted to capture the risk factors associated with the zoonoses under study. Household was considered as a cluster. In addition to individuals within the selected households, people from the village who fulfilled the criteria (being ≥ 16 years, whose animals sampled and had willingness to participate the study) were voluntarily selected and sampled. A total of 190 humans were sampled from six kebeles. All the samples (n=190) were used for ELISA test but only 178 were used for brucellosis screening using RBPT.

### 2.4. Questionnaire administration

Households whose livestock and/people were sampled were questioned about livestock health and management as well as people demographic information and their risky practices. Some of the information was used to analyse the risk factors. The questionnaire was translated from English to Somali.

### 2.5. Blood samples collection

A nurse collected blood samples by venipuncture in 5 ml vacutainer tubes from humans and a veterinarian used 10 ml plain vacutainer tubes for livestock. The blood samples were labeled and kept at room temperature until clot formation. The blood samples were centrifuged at 3000 rpm for 5 minutes. Sera were separated using pasteur pipettes and placed in a labeled 2 ml Eppendorf sera tubes. Sera samples were transported on ice to Gode city and stored at −20°C until transported to Addis Ababa for laboratory testing at the Armauer Hansen Research Institute.

### 2.6. Serological tests

#### 2.6.1. Brucellosis serology

Sera samples were first screened with the RBPT (ID. vet, Innovative Diagnostics, RSA-RB ver 0112 GB, Grabes, France). In livestock, all samples (n=872) were screened by RBPT but only 141 camels, 252 goats, 229 sheep and 108 cattle (n=730) were then further tested by ELISA test. The reagents were left under room temperature for 30 minutes before testing. Equal volume of the reagent and serum (30μl) were placed on a clean plate. First, 30 μl of Rose Bengal was placed on the plate and 30 μl of serum was added then mixed thoroughly by using wooden applicator sticks and then the plate was shaken slowly with hand for about 4 minutes [44]. Any visible agglutination by naked eyes was considered as positive and lack of agglutination was considered as negative. Even if slight agglutination was observed, it was considered as a positive. Human sera which were positive in RBT (n=5) were sequentially diluted with phosphate buffered saline (PBS) to obtain dilutions from1/4 and 1/8. All sera were found reactive in 1/4 dilutions and three sera were also reactive in 1/8 dilution.

All livestock samples positive with the RBPT(n=23) were further tested by indirect ELISA (CHEKIT Brucellose Serum ELISA Test Kit, IDEXX Laboratories, ME, USA) and classified as positive or negative according to the manufacturer’s recommended cut-off ranges. Samples were tested in duplicates and the mean optical density (OD) value at 450nm of each was calculated [(S_ample_/P_ositive_% = mean OD sample – mean OD negative control/ (mean OD positive control – mean OD NC) x100]. Brucellosis results were interpreted as positive (S/P ≥ 80%) and negative (S/P < 80%). Results were checked for validity according to the manufacturer’s recommendations. In livestock, only iELISA positive samples were used for the data analysis, whereas in humans, RBPT positive samples were used for the data analysis.

#### 2.6.2. Q-fever and Rift Valley Fever serology

All ruminants and camels samples were tested using indirect ELISA for Q-fever by using *Coxiella burnetii* phase I and II strain (ID-vet, Innovative Diagnostics, FQS-MS ver 0514 GB, Grabes, France). The Panbio *Coxiella burnetii* (Q-Fever) IgG ELISA was used for human sera (Panbio diagnostics, Cat. no. 06PE10, Germany). Q-fever results of livestock were classified as seropositive and seronegative by calculating the S/P% as described above. Q-fever results of livestock were interpreted as positive (S/P > 50%) and negative (S/P ≤ 40%). Q-fever results in humans were interpreted using an index value (IV) (IV= sample absorbance/cut-off value) as positive (IV > 1.1) and negative (IV < 0.9). All equivocal (doubtful) human Q-fever samples were re-tested. Results were checked for validity according to the manufacturer’s recommendations.

Competitive ELISA (ID-vet, Innovative Diagnostics, RIFTC ver 1114 GB, Grabes, France) was used for Rift Valley Fever in both humans and livestock. RVF results were classified as seropositive and seronegative by calculating the mean OD value of each sample in both humans and livestock. Results were expressed as percentage (S_ample_ /N_egative_ % = OD _sample_ /OD _NC_ × 100) and interpreted as positive (S/N ≤ 40%) and negative (S/N > 50%).

### 2.7. Data analysis

The data was entered into Microsoft Access then analyzed using STATA version 14 (Stata Corporation, College Station, TX, USA). Both descriptive and analytical statistics were used for data analysis. Logistic regression with clustering at household/herd level was used to estimate the apparent seroprevalence of humans and livestock. Uni and multivariable analysis was done to identify predictors for seropositivity. Age category, sex and kebele were included as categorical variables in the pre specified multivariable model. Age categories varies according to species. For sheep and goats (young= 1-2 years, adult= 3-6 and old= >6). For cattle (young= 1-3 years, adult= 4-7 and old= >7). For camels (young= 1-4 years, adult= 5-8 and old= >8). For humans (young adult= 16-31 years, middle-aged adult= 32-48 and old adult= ≥ 49). Generalized Estimating Equation (GEE) model for binomial outcomes were used to account for potential correlation within herds. For the correlation matrix in figure 3, we calculated the pairwise Pearson’s correlation coefficient for the prevalence in two different species.

### 2.8. Ethical clearance

The study received ethical clearance from the “Ethikkommission Nordwest-und Zentralschweiz” (EKNZ) in Switzerland (BASEC UBE-req.2016-00204) and the Jigjiga University Research Ethics Review Committee (JJU-RERC002/2016).

## 3. Results

### 3.1. Descriptive analysis of the study population

About 77.4% (565/730) of the livestock were females and 22.6% (165/730) were males. About half of livestock sampled were adults; cattle (49.1%), camels (45.4%), goats (61.1%) and sheep (0%). In human samples, 48.9% (93/190) were females and 51.1% (97/190) were males with mean age of 42 years. The mentioned zoonotic diseases by the respondents included brucellosis, tuberculosis, and anthrax.

The livestock vaccination status was based on all types of vaccines provided by the government except those against zoonotic diseases under the study (Table 1).

**Table 1.** Sampled household related information

| Variable | Category | (% or mean±SD<sup>a</sup>) |
| --- | --- | --- |
| Family size | 1-6 | 25 |
|  | 7-10 | 54 |
| | $\geq 11$ | 21 |
| Production system | Agropastoral | 38 |
|  | Pastoral | 62 |
| Livestock disease event prior to | Abortion | 90 |
|  | Retained placenta | 38 |
| 6 months | Weak newborns | 60 |
| Livestock vaccination status | Non-vaccinated | 41 |
|  | Vaccinated | 59 |
|  | Cattle | 2.0±1.2 |
| Family herd size | Camel | 1.6±1.1 |
|  | Goat | 3.5±1.0 |
|  | Sheep | 3.4±1.0 |
| Milk consumption habit | Raw | 87 |
|  | Boiled | 13 |
| Zoonoses mentioned among all reported herd diseases | Mentioned at least one | 10 |
|  | Mentioned as zoonoses but were not zoonoses | 17 |
|  | I do not know | 73 |
|  | Married | 99 |
| Marital status | Single | 1 |
| Zoonoses awareness | Yes | 27 |
|  | No | 73 |
| Animal delivery assistance | Yes | 100 |
|  | No | 0 |
| Aborted fetus disposal | Throw in the field | 100 |
|  | Burn | 0 |
|  | Bury | 0 |
|  | Others | 0 |

<sup>a</sup> SD= Standard deviation

### 3.2. Apparent seroprevalence estimates of Q-fever, RVF and brucellosis in humans and livestock in Adadle, Somali region of Ethiopia

The apparent seroprevalence of Q-fever in humans was 27.0% (95% CI 20.4-34.0) and RVF was 13.2% (95% CI 8.7-18.8) (table 2). The apparent seroprevalence of Q-fever and RVF in livestock was 39.0% (95% CI 35.1-42.3) and 15.2% (95% CI 12.7-18.0) respectively. The apparent seroprevalence of brucellosis in humans was 2.8% (0.9-6.4) and 1.5% (0.2-5.2), 0.6% (0.0-3.2) in cattle and camels respectively (table 2).

**Table 2.** Apparent seroprevalence of Q-fever, RVF and brucellosis in humans and livestock

| Zoonoses | Species | n-tested | n pos | Apparent (95% CI<sup>a</sup>) |
| --- | --- | --- | --- | --- |
| Q-fever | Human | 188 | 50 | 26.3(20.2-33.4) |
|  | Cattle | 108 | 11 | 9.6 (5.9-15.1) |
|  | Camel | 141 | 79 | 55.7(46.0-65.0) |
|  | Goat | 252 | 123 | 48.8(42.5-55.0) |
|  | Sheep | 229 | 69 | 28.9(25.0-33.2) |
| RVF | Human | 190 | 25 | 13.2(8.7-19.4) |
|  | Cattle | 108 | 19 | 17.9(11.0-27.8) |
|  | Camel | 141 | 60 | 42.6(34.8-50.7) |
|  | Goat | 252 | 15 | 6.3(3.3-11.6) |
|  | Sheep | 229 | 17 | 7.4(4.7-11.5) |
| Brucellosis | Human | 178 | 5 | 2.8(1.2-6.5) |
|  | Cattle | 135 | 2 | 1.5 (0.4-5.6) |
|  | Camel | 171 | 1 | 0.6(0.1-4.0) |
|  | Goat | 297 | 0 | -- |
|  | Sheep | 269 | 0 | -- |

<sup>a</sup> 95% CI are adjusted for clustering

In livestock, the highest seroprevalence of Q-fever was found in Harsog (50.0%, 95% CI 41.4-58.6) and the least in Higlo (29.1%, 95% CI 17.6-42.9). In humans, the highest seroprevalence of Q-fever was recorded in Boholhagare (42.0%, 95% CI 28.2-57.0) and the least in Gabal (5.9%, 95% CI 0.1-28.7). The highest seroprevalence of RVF in livestock was found in Bursaredo (19.6%, 95% CI 13.7-26.7) and the least in Melkasalah (9.8%, 95% CI 4.3-18.3). The highest seroprevalence of RVF in humans was 27.5% (95% CI 15.9-41.7) and the least was 4.4% (95% CI 0.5-14.8) in Boholhagare and Harsog respectively (Fig 2).

**Fig 2.**
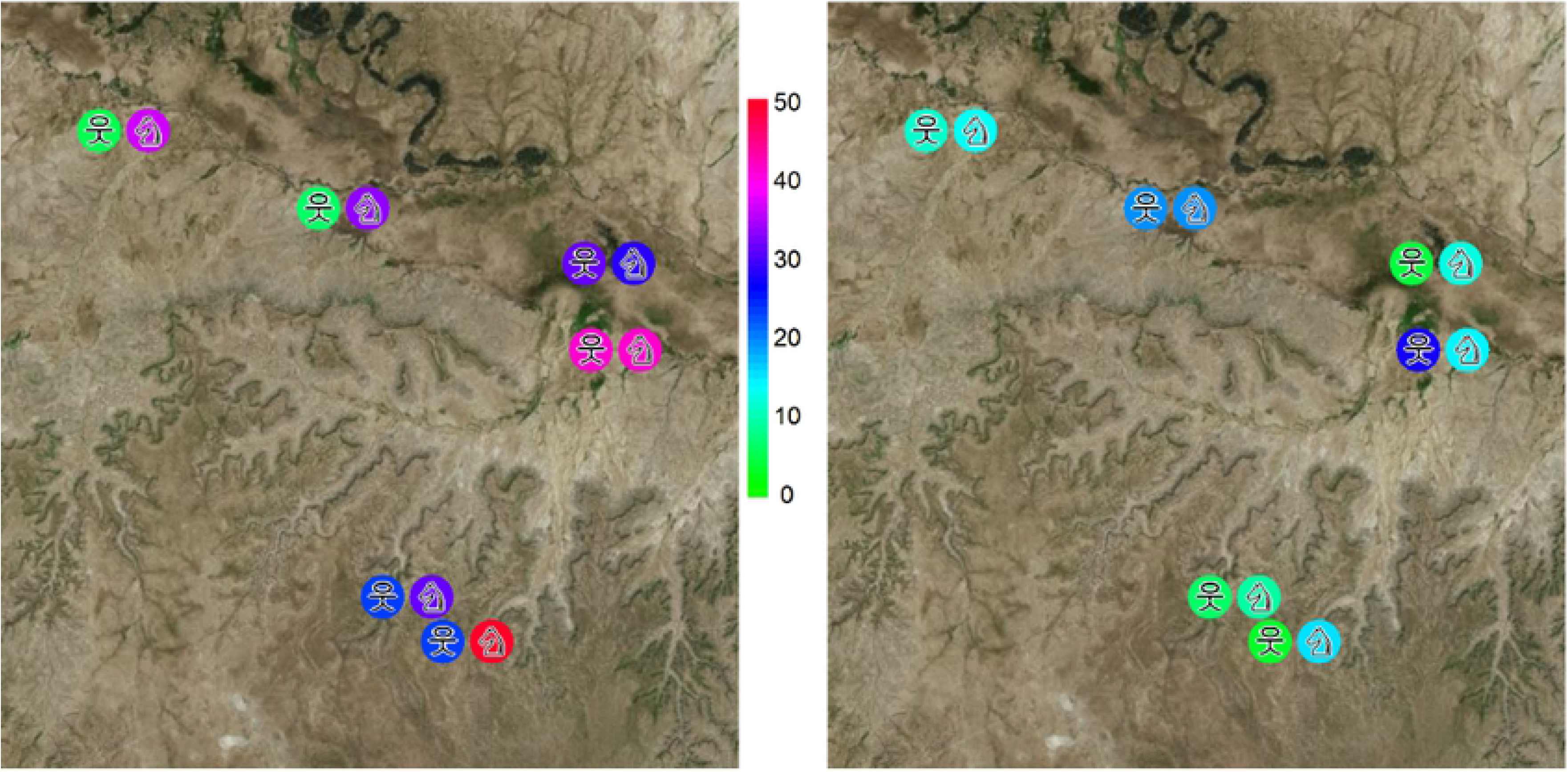
The apparent seroprevalence of Q-fever (left) and RVF (right) in humans and livestock in Adadle woreda, Somali region. 웃 = humans and ♘= livestock.

Camels had the highest seroprevalence of both Q-fever and RVF at herd level with 55.7% (95% CI 46.0-65.0) and 42.6% (95% CI 34.8-50.7) respectively. The lowest seroprevalence of Q-fever at herd level was found in cattle with 9.6% (95% CI 5.9-15.1) and RVF in goats with 6.3% (95% CI 3.3-11.6) (table 2).

### 3.3. Apparent seroprevalence estimates of brucellosis in humans and livestock in Adadle, Somali region of Ethiopia

The apparent seroprevalence of brucellosis in humans was 2.8% (0.9-6.4) and 0.3% (0.0-1.0) in livestock. Only cattle and camels were found seropositive for iELISA and all were females. The individual seroprevalence was 1.5% (95% CI 0.2-5.2) in cattle and 0.6% (95% CI 0.0-3.2) in camels. Seropositive cattle were from Boholhagare and Gabal kebeles whereas seropositive camels were only from Melkasalah kebele. No correlation was found between risk factors and brucellosis seropositivity in both humans and livestock. All seropositive samples were males in humans and females in livestock. Seropositivity of brucellosis was decreasing as age increased in humans but increased as age increased in cattle. The only positive sample for camel was in the age between five and eight years.

### 3.4. Risk factors associated with human seropositivity of Q-fever and RVF

In contrast to livestock, human seroprevalence was higher in males than females. Males had on average of 30% and 90% odds of seropositivity for Q-fever (OR= 1.3; 95% CI 0.6-2.5) and RVF (OR= 1.9; 95% CI 0.7-4.8) than females respectively. Human seroprevalence increased with increasing age for RVF but not for Q-fever. In multivariable analysis, there were no significant association between any risk factor variables and seropositivity of Q-fever and RVF in humans next to kebele (table 3).

**Table 3.** Risk factors associated with human seropositivity for Q-fever and RVF

|  |  |  | **Q-fever** | Odds ratio (95% CI) |  |  | **RVF** | Odds ratio (95% CI) |  |
| --- | --- | --- | --- | --- | --- | --- | --- | --- | --- |
| Predictors | *Category* | *N tested* | *Number (% seropositive)* | Univariable analysis | Multivariable analysis | *N tested* | *Number (% seropositive)* | Univariable analysis | Multivariable analysis |
| Kebele | Boholhagare | 50 | 21(42.0) | 1 | 1 | 51 | 14(27.5) | 1 | 1 |
|  | Gabal | 17 | 1(5.9) | 0.1(0.0,0.7) | 0.1(0.0,0.7) | 17 | 2(12.0) | 0.4(0.1,1.7) | 0.4(0.1,1.9) |
|  | Harsog | 46 | 11(24.0) | 0.4(0.2,1.0) | 0.5(0.2,1.2) | 46 | 2(4.4) | 0.1(0.0,0.6) | 0.1(0.0,0.7) |
|  | Higlo | 19 | 6(32.0) | 0.6(0.2,2.0) | 0.7(0.2,2.1) | 19 | 1(5.3) | 0.1(0.0,1.2) | 0.2(0.0,1.3) |
|  | Melkasalah | 41 | 10(24.4) | 0.4(0.2,1.1) | 0.5(0.2,1.1) | 42 | 3(7.1) | 0.2(0.1,0.8) | 0.2(0.1,0.9) |
|  | Bursaredo | 15 | 1(6.7) | 0.1(0.0,0.8) | 0.1(0.0,0.8) | 15 | 3(20.0) | 0.7(0.2,2.7) | 0.6(0.1,2.6) |
| Sex | Female | 91 | 22(24.2) | 1 | 1 | 93 | 8(9.0) | 1 | 1 |
|  | Male | 97 | 28(28.9) | 1.2(0.6,2.3) | 1.3(0.6,2.5) | 97 | 17(18.0) | 2.2(0.9,5.4) | 1.9(0.7,4.8) |
| Age | 16-31 | 68 | 17(25.0) | 1 | 1 | 85 | 8(9.4) | 1 | 1 |
|  | 32-48 | 55 | 13(24.0) | 1.0(0.4,2.1) | 1.0(0.4,2.2) | 40 | 5(13.0) | 1.2(0.4,4.0) | 1.0(0.3,3.6) |
|  | ≥49 | 65 | 20(31.0) | 1.3(0.6,2.8) | 1.3(0.6,3.1) | 65 | 12(18.5) | 1.9(0.7,4.8) | 1.5(0.5,4.3) |

### 3.5. Risk factors associated with livestock seropositivity for Q-fever and RVF

In livestock, high seroprevalence of both diseases were found in female animals than males and older age animals (except sheep). In sheep, all seropositive samples were older than six years. The cattle with age 4-7 years had higher odds of getting Q-fever infection than those less than 4 years (OR= 2.5; 95% CI 0.2-29.6) but the confidence interval was broad and included unity. Camel seropositivity of Q-fever and RVF were significantly associated with age (OR= 8.1; 95% CI 2.8-23.7 and OR=8.4; 95% CI 2.3-30.3) respectively (Table 4).

**Table 4.**
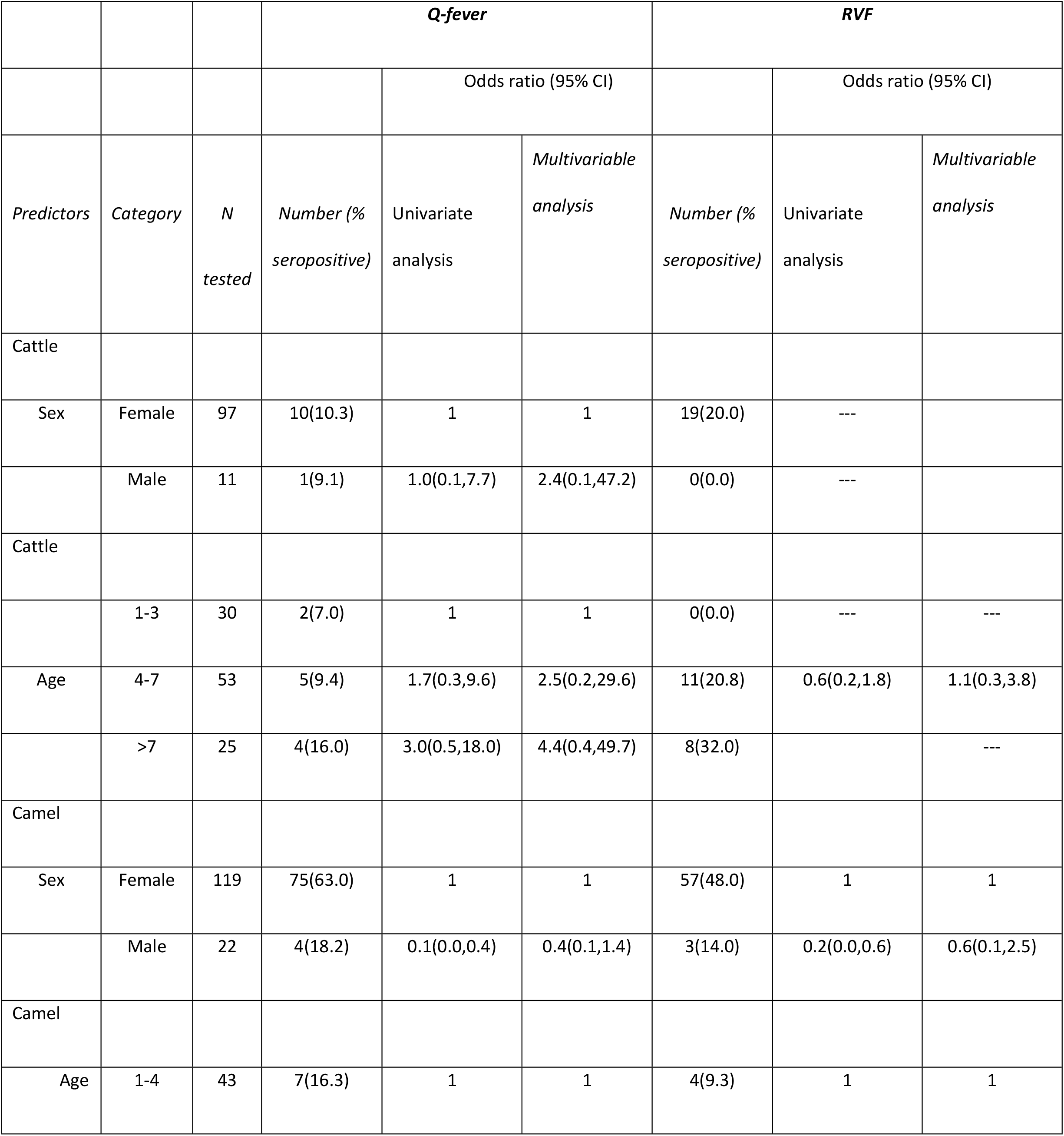
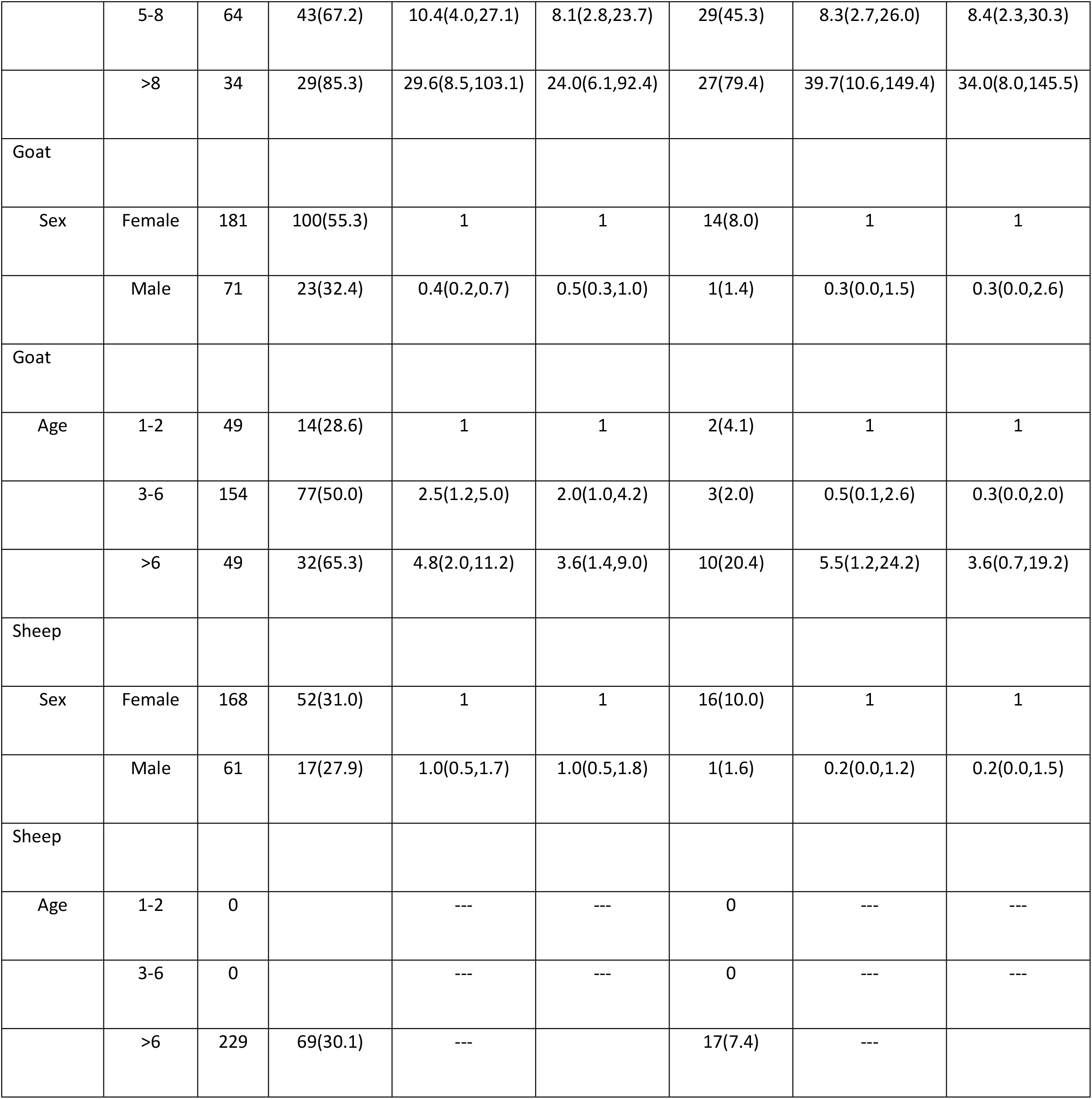
Risk factors associated with livestock seropositivity for Q-fever and RVF

|  |  |  | *Q-fever* |  |  | *RVF* |  |  |
| --- | --- | --- | --- | --- | --- | --- | --- | --- |
|  |  |  |  | Odds ratio (95% CI) |  |  | Odds ratio (95% CI) |  |
| *Predictors* | *Category* | *N tested* | *Number (% seropositive)* | Univariate analysis | *Multivariable analysis* | *Number (% seropositive)* | Univariate analysis | *Multivariable analysis* |
| Cattle |  |  |  |  |  |  |  |  |
| Sex | Female | 97 | 10(10.3) | 1 | 1 | 19(20.0) | --- |  |
|  | Male | 11 | 1(9.1) | 1.0(0.1,7.7) | 2.4(0.1,47.2) | 0(0.0) | --- |  |
| Cattle |  |  |  |  |  |  |  |  |
|  | 1-3 | 30 | 2(7.0) | 1 | 1 | 0(0.0) | --- | --- |
| Age | 4-7 | 53 | 5(9.4) | 1.7(0.3,9.6) | 2.5(0.2,29.6) | 11(20.8) | 0.6(0.2,1.8) | 1.1(0.3,3.8) |
|  | >7 | 25 | 4(16.0) | 3.0(0.5,18.0) | 4.4(0.4,49.7) | 8(32.0) |  | --- |
| Camel |  |  |  |  |  |  |  |  |
| Sex | Female | 119 | 75(63.0) | 1 | 1 | 57(48.0) | 1 | 1 |
|  | Male | 22 | 4(18.2) | 0.1(0.0,0.4) | 0.4(0.1,1.4) | 3(14.0) | 0.2(0.0,0.6) | 0.6(0.1,2.5) |
| Camel |  |  |  |  |  |  |  |  |
| Age | 1-4 | 43 | 7(16.3) | 1 | 1 | 4(9.3) | 1 | 1 |
|  | 5-8 | 64 | 43(67.2) | 10.4(4.0,27.1) | 8.1(2.8,23.7) | 29(45.3) | 8.3(2.7,26.0) | 8.4(2.3,30.3) |
|  | >8 | 34 | 29(85.3) | 29.6(8.5,103.1) | 24.0(6.1,92.4) | 27(79.4) | 39.7(10.6,149.4) | 34.0(8.0,145.5) |
| Goat |  |  |  |  |  |  |  |  |
| Sex | Female | 181 | 100(55.3) | 1 | 1 | 14(8.0) | 1 | 1 |
|  | Male | 71 | 23(32.4) | 0.4(0.2,0.7) | 0.5(0.3,1.0) | 1(1.4) | 0.3(0.0,1.5) | 0.3(0.0,2.6) |
| Goat |  |  |  |  |  |  |  |  |
| Age | 1-2 | 49 | 14(28.6) | 1 | 1 | 2(4.1) | 1 | 1 |
|  | 3-6 | 154 | 77(50.0) | 2.5(1.2,5.0) | 2.0(1.0,4.2) | 3(2.0) | 0.5(0.1,2.6) | 0.3(0.0,2.0) |
|  | >6 | 49 | 32(65.3) | 4.8(2.0,11.2) | 3.6(1.4,9.0) | 10(20.4) | 5.5(1.2,24.2) | 3.6(0.7,19.2) |
| Sheep |  |  |  |  |  |  |  |  |
| Sex | Female | 168 | 52(31.0) | 1 | 1 | 16(10.0) | 1 | 1 |
|  | Male | 61 | 17(27.9) | 1.0(0.5,1.7) | 1.0(0.5,1.8) | 1(1.6) | 0.2(0.0,1.2) | 0.2(0.0,1.5) |
| Sheep |  |  |  |  |  |  |  |  |
| Age | 1-2 | 0 |  | --- | --- | 0 | --- | --- |
|  | 3-6 | 0 |  | --- | --- | 0 | --- | --- |
|  | >6 | 229 | 69(30.1) | --- |  | 17(7.4) | --- |  |

### 3.6. Correlation between human seropositivity and livestock seropositivity for Q-fever and RVF

Generally, there was only a weak correlation between human seropositivity and livestock seropositivity for both Q-fever and RVF. Human seropositivity of Q-fever was related with goats and RVF seropositivity was related with camels (Fig 3).

**Fig 3.**
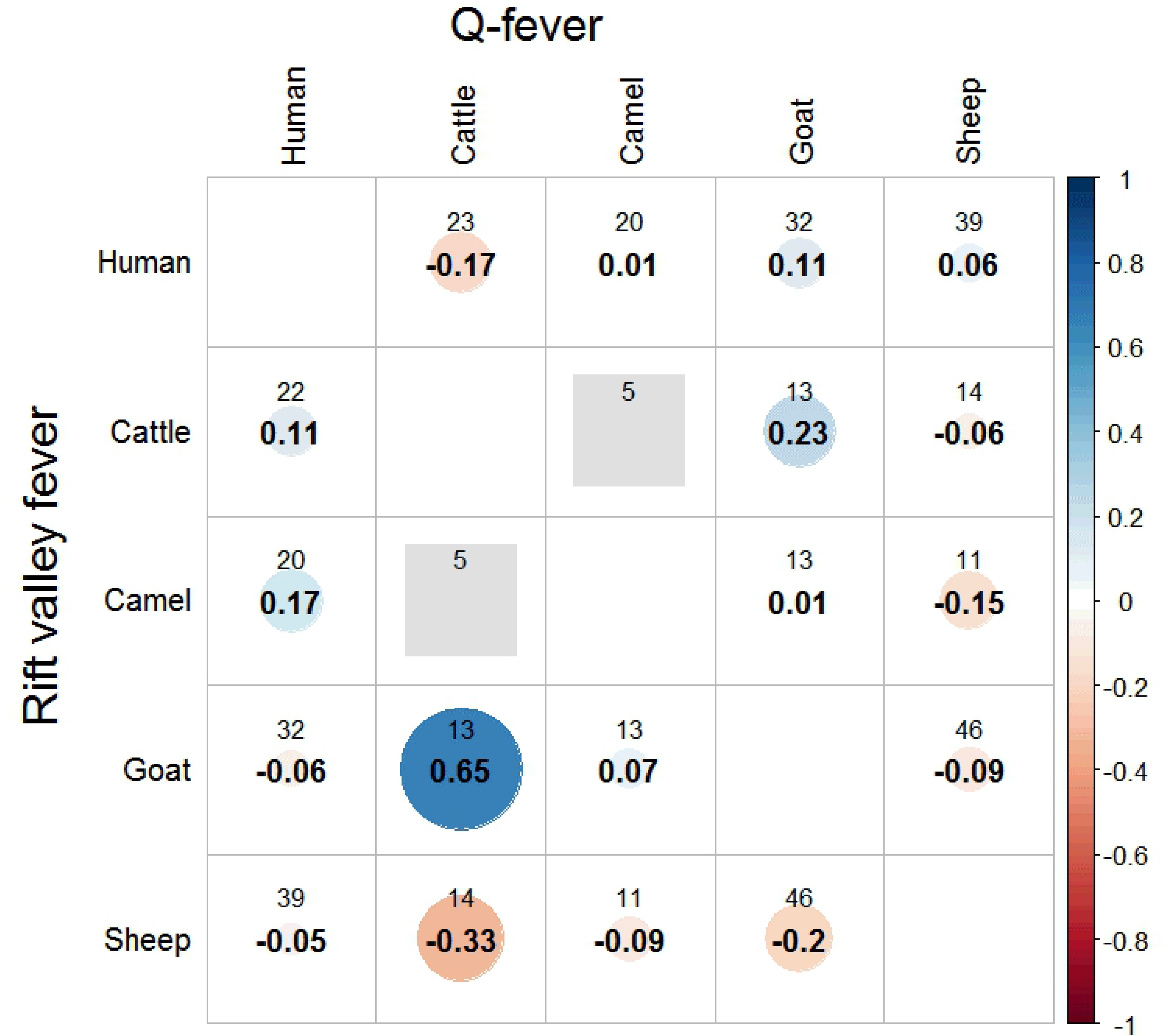
Correlation between humans and livestock seropositivity for Q-fever and RVF. The upper number shows herd number and the lower number shows the Pearson’s correlation coefficient.

## 4. Discussion

The current findings established the seroprevalence of brucellosis, Q-fever and RVF in humans and livestock using for the first time a One Health study approach in the Somali region of Ethiopia. Mainly female animals were found in the sampled households, since pastoral communities keep animals mainly for reproduction and milk purposes. Agropastoral kebeles mostly kept small ruminants and cattle whereas in pastoral kebeles, they kept camels and small ruminants. Pastoralists had a nomadic way of life whereas agropastoralists were either transhumant or settled. Livestock abortions (90%) and weak newborns (60%) were commonly reported (Ibrahim et al., in press) and might cause negative consequences in production and economy for the households. According to our study, brucellosis was not the causative agent for abortion. There might be other infectious or non-infectious diseases causes that needs to be researched in the future. Camel abortion outbreak occurred in Somali region in 2016, and all samples tested found negative for brucellosis (Muhumed Ali, SORPARI staff; personal communication). Information about abortion incidences of pastoral livestock in Ethiopia that are vastly kept in the low lands are lacking. Abortion incidences in Ethiopia dairy cows in the highlands ranged from 2.2%- 28.9% [45].

This current finding of brucellosis seroprevalence was low. This was comparable with previous studies [11, 46] in camels and [11, 47-49] in cattle which reported from Somali and Oromia regions of Ethiopia. However, this study showed a lower prevalence than other previous studies in Ethiopia [12, 50, 51]. This difference might be due to variation in location, husbandry and management system, breed and type of serological tests used [47, 52]. Most of the studies conducted in Ethiopia were used complement fixation test as confirmatory diagnosis unlike the current study, which used iELISA. All small ruminants (n=11) which were seropositive in RBT were seronegative in iELISA. This might be that more false positives were captured by RBT but were seronegative using iELISA. Similarly [2, 20] found 0% seroprevalence in small ruminants in Cote d’Ivoire and Togo. All seropositive were males in humans and females in livestock. Seropositivity of brucellosis in only female livestock shows their susceptibility for the infection and dominance within the herd [50]. The seropositivity of brucellosis had decreased as age increased in humans but increased as age increased in cattle. Higher seropositivity in older ages might be due to high risk of infection because of age and the multiple parities as they got older [53].

Q-fever studies in Ethiopia are rare and the few available studies focused on ticks. The present findings confirmed high Q-fever seroprevalence in humans and livestock. This is in agreement with the study [54]. Camels had the highest seroprevalence for both Q-fever and RVF. Highest Q-fever seropositivity in camels was in agreement with a study from Gumi et al., (2013) in southern pastoralist livestock of Ethiopia. The seroprevalence found in camels was lower than the above cited study, which might be due to differences in the study locations [55], however, was comparable to other studies [21, 56]. Previous studies in Ethiopia showed that seroprevalence of brucellosis were lower in eastern than southern parts of the country which could hold true for Q-fever too [14, 57]. Relatively higher Q-fever seroprevalence in both humans and livestock were recorded in agropastoral than in pastoral kebeles.

Tick infestation was reported to be higher in agropastoral than pastoral kebeles (Ibrahim et al., in press). Ticks are naturally infected by *Coxiella burnetii* and transmit the *Coxiella* from infected animals during their blood meal to other healthy animals. We have observed that the communities used ineffective diazinone as acaricide indicating that ticks were regarded by enrolled communities as a livestock health problem (Ibrahim et al., in press). The diazinone was not effective either because it was available in the market informally through from Somalia where the quality was poor as compared to the ones imported formally into the country or pastoralists used it themselves with sometimes inappropriate dilution concentration.

In agropastoral kebeles, high wind movements were observed during the dry season (June-August). Human Q-fever infection are likely to occur where livestock seroprevalence is high and such winds are common facilitating the inhalation of dust contaminated with *Coxiella* that are spread massively by livestock during abortions due to Q-fever [58]. It was common in the area to assist animal delivery with bare hands and inappropriate management of aborted fetus, which could increase the exposure of the disease [59]. In our study, human Q-fever seropositivity was weakly correlated with goats. This is in contrast to previous studies [23, 55, 60], but in line with recent outbreaks in Canada, Australia and Netherlands [18, 61, 62].

Seroprevalence of Q-fever in female camels were three times higher than males. The same pattern was observed among other livestock species. Similar findings were found in various studies in the Sahel [56, 63]. This might be due to high susceptibility of the bacteria to udder, placenta and amniotic fluids. Seroprevalence of Q-fever in camels was statistically significant associated with age (p<0.001). This was comparable with the study of [63]. Another studies showed that, like in our study-increasing age increased the seroprevalence of Q-fever in all livestock species [64–66] which is not surprising given the cumulative time of potential exposure. Unlike livestock, men had twice higher seroprevalence for Q-fever than women. This might be that, males took livestock to the market and are exposed to contaminated dusts (Ibrahim et al., in press).

There has been recently an increasing evidence and documentation of RVF inter-epidemic cases in East and central Africa [34–37]. To our knowledge, this study is the first to report RVF seropositivity in humans and livestock in Ethiopia. Different models predicted the suitability of RVF occurrence in Ethiopia due to climate change, vector distribution and livestock exchanges with neighboring countries with history of RVF outbreaks [33, 38]. This study showed high seroprevalence of RVF in both humans and livestock, which lay within the ranges of reported seroprevalences in other East African countries [26]. For livestock, relatively high seroprevalences of RVF were found in agropastoral kebeles for camels and cattle, but these were not significantly different to those of small ruminants. High human seroprevalence of RVF was found in our study in agropastoral kebeles. This could be due to the abundance of vectors in those kebeles closer to the river (1-18 km) and main livestock species (sheep and goats) susceptibility for RVF-virus. Flooding of the Wabi-Shabele river is common in these agropastoral kebeles of Adadle woreda which might increase the suitability of amplification and transmission of RVF-Virus similar to the report by [67] in Madagascar. In contrast to our study, Sumaye et al., (2013) reported high seroprevalence the further away from flooding area in Tanzania.

Agropastoral kebeles were relatively nearer than pastoral kebeles to the largest livestock market (Gode) in the area. At Gode market, animals from different areas including neighboring Somalia are traded. Hence, high livestock movements for trade might increase RVFV exposure [35]. RVF seropositivity was associated with livestock species. Among all livestock species, seroprevalence of RVF was statistically significant with increasing age only in camels. Traditionally in pastoral communities, camels were rarely sold especially females compared to other livestock species. This might increase the exposure of RVFV in female camels as they stay long in the herd. Indeed, it also shows RVFV exposure in the area since a longer period. What seems important to highlight is the fact that in small ruminants and camels we found seropositivity also in the youngest class, which suggests ongoing (inter-epidemic) transmission. The risk of human exposure during inter-epidemic livestock infection is not yet well documented. However, one can state that an endemic situation on livestock most likely leads to endemic infection pressure in people. Unlike livestock, men had twice higher seroprevalence for RVF than women. This was similar with the study of [68]. Human seropositivity for RVF increased with age. This might be the potential risk of older people to be exposed to infected materials and vector for RVFV as in Kenya [37] or the longer you live, the higher chance to get once in your life exposure to the agent.

Assessing human and livestock zoonoses seroprevalence simultaneously allowed the identification of the most important animal sources. In this way, an added value of an integrated human and animal health approach is demonstrated. More researches is needed to use this data in view of using it to plan cost-effective intervention programs-and then to compare to other human and animal health priorities.

## Conclusions

This study revealed the exposure to brucellosis, Q-fever, and RVF in humans and livestock in Adadle woreda. Our results indicated that there are several zoonotic infections in the area without clinical signs or outbreaks. The medical personnel should consider such zoonoses more carefully because most cases were either misreported or ignored at all in the daily routine diagnosis at health facilities. Hence, continuous sero-surveillance in both humans and livestock is necessary. Further researches to look more in depth into negotiating health priorities and intervention strategies in face of other prevailing health problems in people and livestock is needed. A One Health study approach as used here allowed to detect most important sources for people of three zoonotic diseases and provided evidence of needed future negotiations on potential actions in surveillance and interventions.

## 5. Acknowledgments

We would like to thank the Swiss Agency for Development and Cooperation (SDC) for funding this study. This study would not have been possible without the support of Adadle communities especially, woreda administrators, human and animal health bureaus, zonal administrators and pastoral/agropastoral communities. We would like to thank the Armauer Hansen Research Institute (AHRI) for the support with the lab analysis, particularly Robel Gesese, Azeb Tarekegn, AshenafiG/giorgis, Ashenafi Alemu, Mahlet Usman, Marechign Yimer, Metasebiya Tegegn, Melaku Tilahun and Biruk Yeshitela. Lastly, we would like to thank Jigjiga University and the JOHI team for their collaboration and support.

## References

1. World Bank. People, Pathogens and our Planet. The Economics of One Health. 2012. Report No.: 69145–GLB.

2. Kanoute YB, Gragnon BG, Schindler C, Bonfoh B, Schelling E. Reprint of “Epidemiology of brucellosis, Q Fever and Rift Valley Fever at the human and livestock interface in northern Cote d’Ivoire”. Acta Trop. 2017;175:121–9.

3. Zinsstag J, Schelling E, Waltner-Toews D, Whittaker M, Tanner M. One Health: the theory and practice of integrated health approaches: CABI; 2015.

4. Grace D, Mutua F, Ochungo P, Kruska R, Jones K, Brierley L, et al. Mapping of poverty and likely zoonoses hotspots. 2012.

5. Pieracci EG, Hall AJ, Gharpure R, Haile A, Walelign E, Deressa A, et al. Prioritizing zoonotic diseases in Ethiopia using a one health approach. One Health. 2016;2:131–5.

6. McDermott John, Grace Delia, Zinsstag J. Economics of brucellosis impact and control in low-income countries. 2013;32(1):249–61.

7. Ducrotoy MJ, Ammary K, Ait Lbacha H, Zouagui Z, Mick V, Prevost L, et al. Narrative overview of animal and human brucellosis in Morocco: intensification of livestock production as a driver for emergence? Infect Dis Poverty. 2015;4:57.

8. Al Shehhi N, Aziz F, Al Hosani F, Aden B, Blair I. Human brucellosis in the Emirate of Abu Dhabi, United Arab Emirates, 2010-2015. BMC Infect Dis. 2016;16(1):558.

9. Sharma HK, Kotwal SK, Singh DK, Malik MA, Kumar A, Rajagunalan, et al. Seroprevalence of human brucellosis in and around Jammu, India, using different serological tests. Vet World. 2016;9(7):742–6.

10. Garcell HG, Garcia EG, Pueyo PV, Martin IR, Arias AV, Alfonso Serrano RN. Outbreaks of brucellosis related to the consumption of unpasteurized camel milk. J Infect Public Health. 2016;9(4):523–7.

11. Gumi B, Firdessa R, Yamuah L, Sori T, Tolosa T, Aseffa A, et al. Seroprevalence of Brucellosis and Q-Fever in Southeast Ethiopian Pastoral Livestock. J Vet Sci Med Diagn. 2013;2(1).

12. Tilahun B, Bekana M, Belihu K, Zewdu EJJoVM, Health A. Camel brucellosis and management practices in Jijiga and Babile districts, Eastern Ethiopia. 2013;5(3):81–6.

13. Haileselassie M, Kalayou S, Kyule M, Asfaha M, Belihu K. Effect of Brucella infection on reproduction conditions of female breeding cattle and its public health significance in Western tigray, northern ethiopia. Vet Med Int. 2011;2011:354943.

14. Yilma M, Mamo G, Mammo B. Review on Brucellosis Sero-prevalence and Ecology in Livestock and Human Population of Ethiopia. Achievements in the Life Sciences. 2016;10(1):80–6.

15. Yeshwas F, Desalegne M, Gebreyesus M, Mussie HMJEVJ. Study on the seroprevalence of small ruminant brucellosis in and around Bahir Dar, north west Ethiopia. 2011;15(2):35–44.

16. Adugna K, Agga G, Zewde GJRST. Seroepidemiological survey of bovine brucellosis in cattle under a traditional production system in western Ethiopia. 2013;32(3):765–73.

17. Zerfu B, Medhin G, Mamo G, Getahun G, Tschopp R, Legesse M. Community-based prevalence of typhoid fever, typhus, brucellosis and malaria among symptomatic individuals in Afar Region, Ethiopia. PLoS Negl Trop Dis. 2018;12(10):e0006749.

18. Bond KA, Vincent G, Wilks CR, Franklin L, Sutton B, Stenos J, et al. One Health approach to controlling a Q fever outbreak on an Australian goat farm. Epidemiol Infect. 2016;144(6):1129–41.

19. Brandwagt DA, Herremans T, Schneeberger PM, Hackert VH, Hoebe CJ, Paget J, et al. Waning population immunity prior to a large Q fever epidemic in the south of The Netherlands. Epidemiol Infect. 2016;144(13):2866–72.

20. Dean AS, Bonfoh B, Kulo AE, Boukaya GA, Amidou M, Hattendorf J, et al. Epidemiology of brucellosis and q Fever in linked human and animal populations in northern togo. PLoS One. 2013;8(8):e71501.

21. Wardrop NA, Thomas LF, Cook EA, de Glanville WA, Atkinson PM, Wamae CN, et al. The Sero-epidemiology of Coxiella burnetii in Humans and Cattle, Western Kenya: Evidence from a Cross-Sectional Study. PLoS Negl Trop Dis. 2016;10(10):e0005032.

22. Nahed HG, Khaled AJJAS. Seroprevalence of Coxiella burnetii antibodies among farm animals and human contacts in Egypt. 2012;8:619–21.

23. Schelling E, Diguimbaye C, Daoud S, Nicolet J, Boerlin P, Tanner M, et al. Brucellosis and Q-fever seroprevalences of nomadic pastoralists and their livestock in Chad. Prev Vet Med. 2003;61(4):279–93.

24. Abebe A. Prevalence of Q fever infection in the Addis Ababa abattoir. Ethiopian medical journal. 1990;28(3):119–22.

25. OIE. World Organisation for Animal Health 2016 [Available from: https://www.oie.int/fileadmin/Home/eng/Animal_Health_in_the_World/docs/pdf/Disease_cards/RIFT_VALLEY_FEVER.pdf.

26. Clark MHA, Warimwe GM, Di Nardo A, Lyons NA, Gubbins S. Systematic literature review of Rift Valley fever virus seroprevalence in livestock, wildlife and humans in Africa from 1968 to 2016. PLoS Negl Trop Dis. 2018;12(7):e0006627.

27. Nakoune E, Kamgang B, Berthet N, Manirakiza A, Kazanji M. Rift Valley Fever Virus Circulating among Ruminants, Mosquitoes and Humans in the Central African Republic. PLoS Negl Trop Dis. 2016;10(10):e0005082.

28. Ng’ang’a CM, Bukachi SA, Bett BK. Lay perceptions of risk factors for Rift Valley fever in a pastoral community in northeastern Kenya. BMC Public Health. 2016;16:32.

29. Cook EAJ, Grossi-Soyster EN, de Glanville WA, Thomas LF, Kariuki S, Bronsvoort BMC, et al. The sero-epidemiology of Rift Valley fever in people in the Lake Victoria Basin of western Kenya. PLoS Negl Trop Dis. 2017;11(7):e0005731.

30. Abakar MF, Nare NB, Schelling E, Hattendorf J, Alfaroukh IO, Zinsstag J. Seroprevalence of Rift Valley fever, Q fever, and brucellosis in ruminants on the southeastern shore of Lake Chad. Vector Borne Zoonotic Dis. 2014;14(10):757–62.

31. Matiko MK, Salekwa LP, Kasanga CJ, Kimera SI, Evander M, Nyangi WP. Serological evidence of inter-epizootic/inter-epidemic circulation of Rift Valley fever virus in domestic cattle in Kyela and Morogoro, Tanzania. PLoS Negl Trop Dis. 2018;12(11):e0006931.

32. Di Nardo A, Rossi D, Saleh SM, Lejlifa SM, Hamdi SJ, Di Gennaro A, et al. Evidence of Rift Valley fever seroprevalence in the Sahrawi semi-nomadic pastoralist system, Western Sahara. BMC Vet Res. 2014;10:92.

33. Tran A, Trevennec C, Lutwama J, Sserugga J, Gely M, Pittiglio C, et al. Development and Assessment of a Geographic Knowledge-Based Model for Mapping Suitable Areas for Rift Valley Fever Transmission in Eastern Africa. PLoS Negl Trop Dis. 2016;10(9):e0004999.

34. Halawi AD, Saasa N, Pongombo BL, Kajihara M, Chambaro HM, Hity M, et al. Seroprevalence of Rift Valley fever in cattle of smallholder farmers in Kwilu Province in the Democratic Republic of Congo. Trop Anim Health Prod. 2019;51(8):2619–27.

35. Sumaye RD, Geubbels E, Mbeyela E, Berkvens D. Inter-epidemic transmission of Rift Valley fever in livestock in the Kilombero River Valley, Tanzania: a cross-sectional survey. PLoS Negl Trop Dis. 2013;7(8):e2356.

36. Britch SC, Binepal YS, Ruder MG, Kariithi HM, Linthicum KJ, Anyamba A, et al. Rift Valley fever risk map model and seroprevalence in selected wild ungulates and camels from Kenya. PLoS One. 2013;8(6):e66626.

37. LaBeaud AD, Pfeil S, Muiruri S, Dahir S, Sutherland LJ, Traylor Z, et al. Factors associated with severe human Rift Valley fever in Sangailu, Garissa County, Kenya. PLoS Negl Trop Dis. 2015;9(3):e0003548.

38. Anyamba A, Chretien J-P, Small J, Tucker CJ, Formenty PB, Richardson JH, et al. Prediction of a Rift Valley fever outbreak. 2009;106(3):955–9.

39. Gebre-Mariam A. The Critical Issue of Land Ownership. Violent Conflict between the Abdalla Tolo-mogge and the Awlihan in Godey Zone, Somali Region, Ethiopia: NCCR North-South Dialogue; 2007.

40. SRBoFED. Somali Region Bureau of Finance Economic and Development Data Collection Created In ArcGIS 9.3 Using ArcMap. 2014.

41. McDermott JJ, Arimi SJVm. Brucellosis in sub-Saharan Africa: epidemiology, control and impact. 2002;90(1-4):111–34.

42. Bennett S, Woods T, Liyanage WM, Smith DL. A simplified general method for cluster-sample surveys of health in developing countries. 1991.

43. Otte M, Gumm IJPVM. Intra-cluster correlation coefficients of 20 infections calculated from the results of cluster-sample surveys. 1997;31(1-2):147–50.

44. OIE. World Organisation for Animal Health. OIE Terrestrial Manual 2018 [Available from: https://www.oie.int/fileadmin/Home/eng/Health_standards/tahm/3.01.04_BRUCELLOSIS.pdf.

45. Tulu D, Deresa B, Begna F, Gojam AJJoVM, Health A. Review of common causes of abortion in dairy cattle in Ethiopia. 2018;10(1):1–13.

46. Gessese A, Mulate B, Nazir S, Asmare AJJVSMD. Seroprevalence of brucellosis in camels (Camelus dromedaries) in South East Ethiopia. 2014;1:2.

47. Terefe Y, Girma S, Mekonnen N, Asrade B. Brucellosis and associated risk factors in dairy cattle of eastern Ethiopia. Trop Anim Health Prod. 2017;49(3):599–606.

48. Dirar BG, Nasinyama GW, Gelalcha BD. Seroprevalence and risk factors for brucellosis in cattle in selected districts of Jimma zone, Ethiopia. Trop Anim Health Prod. 2015;47(8):1615–9.

49. Degefu H, Mohamud M, Hailemelekot M, Yohannes MJEVJ. Seroprevalence of bovine brucellosis in agro pastoral areas of Jijjiga zone of Somali National Regional State, Eastern Ethiopia. 2011;15(1).

50. Zewdie W. Review on Bovine, Small ruminant and Human Brucellosis in Ethiopia. J Vet Med Res 5(9): 1157. 2018.

51. Bekele WA, Tessema TS, Melaku SKJAvs. Camelus dromedarius brucellosis and its public health associated risks in the Afar National Regional State in northeastern Ethiopia. 2013;55(1):89.

52. Racloz V, Schelling E, Chitnis N, Roth F, Zinsstag JJORSeT. Persistence of brucellosis in pastoral systems. 2013;32(1):61–70.

53. ElTahir Y, Al Toobi AG, Al-Marzooqi W, Mahgoub O, Jay M, Corde Y, et al. Serological, cultural and molecular evidence of Brucella melitensis infection in goats in Al Jabal Al Akhdar, Sultanate of Oman. Vet Med Sci. 2018.

54. Jarelnabi AA, Alshaikh MA, Bakhiet AO, Omer SA, Aljumaah RS, Harkiss GD, et al. Seroprevalence of Q fever in farm animals in Saudi Arabia. 2018;29:895–900.

55. Vanderburg S, Rubach MP, Halliday JE, Cleaveland S, Reddy EA, Crump JA. Epidemiology of Coxiella burnetii infection in Africa: a OneHealth systematic review. PLoS Negl Trop Dis. 2014;8(4):e2787.

56. Hussein MF, Alshaikh MA, Al-Jumaah RS, GarelNabi A, Al-Khalifa I, Mohammed OB. The Arabian camel (Camelus dromedarius) as a major reservoir of Q fever in Saudi Arabia. Comparative Clinical Pathology. 2014;24(4):887–92.

57. Asmare A. Seroprevalence of Brucellosis in Camels (Camelus dromedaries) in South East Ethiopia. Journal of Veterinary Science & Medical Diagnosis. 2014;03(01).

58. Kersh GJ, Wolfe TM, Fitzpatrick KA, Candee AJ, Oliver LD, Patterson NE, et al. Presence of Coxiella burnetii DNA in the environment of the United States, 2006 to 2008. Appl Environ Microbiol. 2010;76(13):4469–75.

59. Tschopp R, Abera B, Sourou SY, Guerne-Bleich E, Aseffa A, Wubete A, et al. Bovine tuberculosis and brucellosis prevalence in cattle from selected milk cooperatives in Arsi zone, Oromia region, Ethiopia. 2013;9(1):163.

60. Abdullah H, El-Shanawany EE, Abdel-Shafy S, Abou-Zeina HAA, Abdel-Rahman EH. Molecular and immunological characterization of Hyalomma dromedarii and Hyalomma excavatum (Acari: Ixodidae) vectors of Q fever in camels. Vet World. 2018;11(8):1109–19.

61. Meadows S, Jones-Bitton A, McEwen SA, Jansen J, Patel SN, Filejski C, et al. Coxiella burnetii (Q Fever) Seropositivity and Associated Risk Factors in Sheep and Goat Farm Workers in Ontario, Canada. Vector Borne Zoonotic Dis. 2016;16(10):643–9.

62. Enserink M. Questions abound in Q-fever explosion in the Netherlands. American Association for the Advancement of Science; 2010.

63. Benaissa MH, Ansel S, Mohamed-Cherif A, Benfodil K, Khelef D, Youngs CR, et al. Seroprevalence and risk factors for *Coxiella burnetii*, the causative agent of Q fever in the dromedary camel (*Camelus dromedarius*) population in Algeria. Onderstepoort J Vet Res. 2017;84(1):e1–e7.

64. Klemmer J, Njeru J, Emam A, El-Sayed A, Moawad AA, Henning K, et al. Q fever in Egypt: Epidemiological survey of Coxiella burnetii specific antibodies in cattle, buffaloes, sheep, goats and camels. PLoS One. 2018;13(2):e0192188.

65. Browne AS, Fevre EM, Kinnaird M, Muloi DM, Wang CA, Larsen PS, et al. Serosurvey of Coxiella burnetii (Q fever) in Dromedary Camels (Camelus dromedarius) in Laikipia County, Kenya. Zoonoses Public Health. 2017;64(7):543–9.

66. Klaasen M, Roest HJ, van der Hoek W, Goossens B, Secka A, Stegeman A. Coxiella burnetii seroprevalence in small ruminants in The Gambia. PLoS One. 2014;9(1):e85424.

67. Chevalier V, Pépin M, Plee L, Lancelot RJES. Rift Valley fever-a threat for Europe? 2010;15(10):18–28.

68. Olive MM, Chevalier V, Grosbois V, Tran A, Andriamandimby SF, Durand B, et al. Integrated Analysis of Environment, Cattle and Human Serological Data: Risks and Mechanisms of Transmission of Rift Valley Fever in Madagascar. PLoS Negl Trop Dis. 2016;10(7):e0004827.

